# Integration of auxin and cytokinin signalling in Arabidopsis root meristem maintenance through the secreted peptide CLE40

**DOI:** 10.1101/2025.07.01.662337

**Authors:** Meik Thiele, Rene Wink, Yvonne Stahl, Madhumitha Narasimhan, Svenja Augustin, Jenia Schlegel, Maike Breiden, Sosan Soltanzadah, Helge Pallakies, Karine Gustavo Pinto, Jonas Schön, Matias D. Zurbriggen, Rüdiger Simon

**Affiliations:** Institute for Developmental Genetics, Heinrich Heine University, Universitätsstr. 1, D-40225 Düsseldorf, Germany; Institute for Molecular Biosciences, Plant Developmental Genetics, Goethe University, Max-von-Laue Str. 9, D-60438 Frankfurt am Main; Miltenyi Biotec B.V. & Co. KG, Friedrich-Ebert-Straße 68, D-51429 Bergisch Gladbach, Germany; QIAGEN GmbH, Qiagen Straße 1, D-40724 Hilden, Germany; Institute for Synthetic Biology, Heinrich Heine University, Universitätsstr. 1, D-40225 Düsseldorf, Germany

## Abstract

Plant meristems perpetually generate new cells for the initiation, growth and differentiation of various cell types and organs. *CLAVATA3/EMBRYO SURROUNDING REGION-RELATED* (*CLE*) genes encode plant specific peptides exerting indispensable functions in a multitude of these developmental processes. Previously, we have described a CLE40 signaling module that regulates stemness in the distal root meristem (DRM). Here, we show that CLE40 is additionally needed to maintain the proximal root meristem (PRM). We demonstrate that in *cle40* mutants, meristematic cells differentiate prematurely, resulting in shortened PRMs and ultimately, reduced root growth. Furthermore, we link these effects to increased cytokinin signaling that can be circumvented by preventing the perception of cytokinin. We highlight how increased cytokinin signaling leads to a change in auxin distribution mediated by *SHORT HYPOCOTYL 2* (*SHY2*), which is ectopically activated if *CLE40* is not active. Finally, we propose that the receptor BARELY ANY MERISTEM 1 (BAM1), but not its homolog BAM2, perceives CLE40 in the PRM.

## Introduction

Plant meristems, which generate the majority of the plant body, are niches that maintain pluripotent stem cells for many years. Stem cells divide slowly and generate more rapidly dividing, transient amplifying (TA) cells and, ultimately, differentiating descendants. The two major stem cell systems, the root and shoot apical meristems (RAM and SAM), are structurally distinct, but share common gene regulatory modules that control their maintenance and adjust cell division rates of the stem cell populations with cell differentiation (**13**,**25**,**55**,**58**). This is achieved through extensive intercellular communication via phytohormones, secreted peptides, mobile transcription factors and small regulatory RNAs. The *Arabidopsis* root stem cell niche (SCN) comprises the quiescent center (QC) consisting of on average four to eight rarely dividing cells, and a surrounding single layer of stem cells or initials that border the QC cells (**17**,**61**). Short range signals that originate from the QC keep the initials in an undifferentiated state. The initials give rise to daughter cells in proximal (towards the shoot), distal (towards the root tip) or lateral directions. The proximal root meristem (PRM), comprising the zone of active cell divisions, increases in size during the first days after germination (DAG) to reach a roughly constant size from 5 DAG onwards (**66**). Towards the apical end of the PRM, cells enter the transition zone (TZ) before undergoing rapid cell elongation and final differentiation. Balanced auxin and cytokinin signaling is critical for PRM homeostasis (**62**).

Secreted peptides of the CLAVATA3/EMBRYO SURROUNDING REGION (CLE) family affect root development in diverse ways (**30**), for example: (1) CLE45 interferes with protophloem development via its receptor BARELY ANY MERISTEM3 (BAM3) (**64**); (2) CLE13 and CLE16 trigger asymmetric cell divisions and generation of new cell layers via the LRR-receptor kinases BAM1/BAM2 and activation of CYCLIND6;1 (**7**); CLE13/CLE16 signalling interacts and partially antagonizes CLE45 signalling in tissue layer organization (**67**); (3) CLE40 restricts the number of columella stem cell layers in the distal root meristem by signaling via the BAM-related CLAVATA1 (CLV1) receptor, and dependent on the presence of ARABIDOPSIS CRINKLY4 (ACR4) (**59**,**60**). Transgenic overexpression of diverse *CLE* genes or exogenous treatment of *Arabidopsis* roots with the corresponding synthetic CLE peptides triggers a rapid shortening of the PRM and premature cellular differentiation (**31**). For these *CLE* functions, the LRR receptor protein CLAVATA2 (CLV2) and the membrane localized receptor kinase CORYNE (CRN) were identified as essential factors, and joint expression of CLV2 with CRN in the protophloem is sufficient to trigger this differentiation response (**26**). Roles for individual CLE peptides in PRM homeostasis have not yet been uncovered, with the exception of CLE40, which not only restricts columella stem cell number, but is also required to support root growth (**28**,**60**). If and how CLE peptide signaling pathways intersect with established hormonal control of root meristem development and maintenance is not entirely understood.

The PLETHORA (PLT) family transcription factors control the position and establishment of the SCN and promote cell division in the PRM in a dose dependent manner (**2**,**5**,**22**,**35**,**57**). *PLT* expression patterns are differentially controlled by an auxin gradient in the root meristem which peaks at the root tip, while PLT protein levels depend on the ROOT MERISTEM GROWTH FACTOR 1 (RGF1)-signalling pathway that interprets ROS (**66**). The distribution of auxin depends on the activities of AUX1-family auxin influx carriers, and on auxin efflux carriers of the PIN-FORMED (PIN) family. Because *PLT* genes promote *PIN* gene expression, they also contribute to the generation of a stable and feedback regulated meristematic region (**22**,**33**,**35**,**56**,**57**). In contrast to auxin signaling which maintains meristematic cell divisions, cytokinin promotes cell differentiation in the TZ via signaling through the transmembrane receptors ARABIDOPSIS HISTIDINE KINASE2 (AHK2), AHK3 and AHK4/WOODEN LEG/CYTOKININ RESPONSE1 (CRE1), which trigger phosphorylation of ARABIDOPSIS RESPONSE REGULATORS (ARRs) (**51**). Type-B ARR transcription factors positively control cytokinin responsive gene expression, while type-A ARRs antagonize this activity (**32**,**51**). In the root meristem, the type-B ARRs ARR1 and ARR12 activate the expression of *SHORT HYPOCOTYL2* (*SHY2*), an AUX/IAA protein that is targeted for degradation in the presence of auxin (**12**,**50**,**51**). *SHY2* is expressed in the vascular tissue of the TZ where it represses the expression of *PIN1,3* and *7,* thereby affecting auxin transport. Loss of *SHY2* leads to an enlarged PRM due to prolonged mitotic activity of TA-cells, whereas auxin-resistant and stabilized *shy2* gain-of-function mutations cause premature consumption of the PRM due to early cell differentiation, resulting in an overall shorter root (**12**,**24**). These studies highlight how the interplay between auxin and cytokinin signaling can act directly and antagonistically determine the overall size of the PRM through *SHY2* (**12**,**38**,**49**).

PRM and SCN development are coordinated through SCARECROW (SCR), a transcription factor which is expressed in the QC, endodermal stem cells and the mature endodermis (**4**,**8**). Besides its role in SCN establishment and cortex/endodermal patterning, SCR also affects *SHY2* expression levels via direct and indirect differential regulation of ARR1: in the QC, SCR represses *ARR1* and cytokinin dependent cell-differentiation; this downregulation of *ARR1* in the QC releases repression of ANTHRANILATE SYNTHASE BETA SUBUNIT1 (ASB1), which controls a key step in auxin biosynthesis; auxin can move from the QC towards the TZ, where it activates *ARR1* expression and thereby *SHY2* to promote cell differentiation (**39**). Thus, genes expressed in the QC determine the overall size of the PRM and the position of the TZ in a non-cellautonomous manner, which is further stabilized by mutual inhibition between PLTs and ARR1 (**51**,**57**). The stability of the PLT2 protein itself depends on the oxidative state within a cell, which is modified by the peptide signal RGF1 that promotes accumulation of superoxide in the meristematic zone (**66**).

Here we show that a loss of CLE40 function results in shorter roots due to premature differentiation of proximal root meristem cells. *CLE40* expression in the procambium is promoted by auxin, and CLE40, by binding to its receptor BAM1, serves to decrease cytokinin levels and signalling. *CLE40* thereby promotes meristem activity and delays differentiation of TA cells. Our findings establish a new function for *CLE40* and uncover how peptide and hormone signaling pathways are tightly interwoven to control root meristem homeostasis and precisely position the transition zone in the root.

## Results

### CLE40 acts in a growth promoting signalling pathway

We explored the role of CLE40 signaling in the regulation of PRM development by comparing the growth behavior of *cle40* mutant roots with that of wild-type plants. *cle40-1* mutants, carrying a transposon insertion in the *CLE40* gene that disrupts the highly conserved CLE motif, were described to develop shorter roots than the corresponding Col-0 wild-type (**28**). We observed the same growth reduction also for the stable *cle40-2* mutant allele (Figure1 A, B, J). Introduction of a transgene expressing an in-frame fusion of the *CLE40* coding region with GFP from the *CLE40* promoter (*pCLE40:CLE40-GFP*) rescued the root length defects of *cle40-2* mutants, thus confirming that the mutation of the *CLE40* gene is causing the growth retardation (**53**). This growth retardation becomes apparent within 4 DAG, and results in a continuous decrease in root growth over time (Figure 1 J). Growth cessation is due to premature differentiation, indicated by a reduced number of normal sized root cortex cells and the resulting decrease of the PRM size, which is only 65 % of wild-type length at 6 DAG in *cle40-2* mutants (Table S1). Root cross sections showed that all cell types are present in *cle40-2* roots, but that the root perimeter was significantly smaller, due to a significant reduction in the number of epidermal and vascular cells (Table S2).

**Figure 1:**
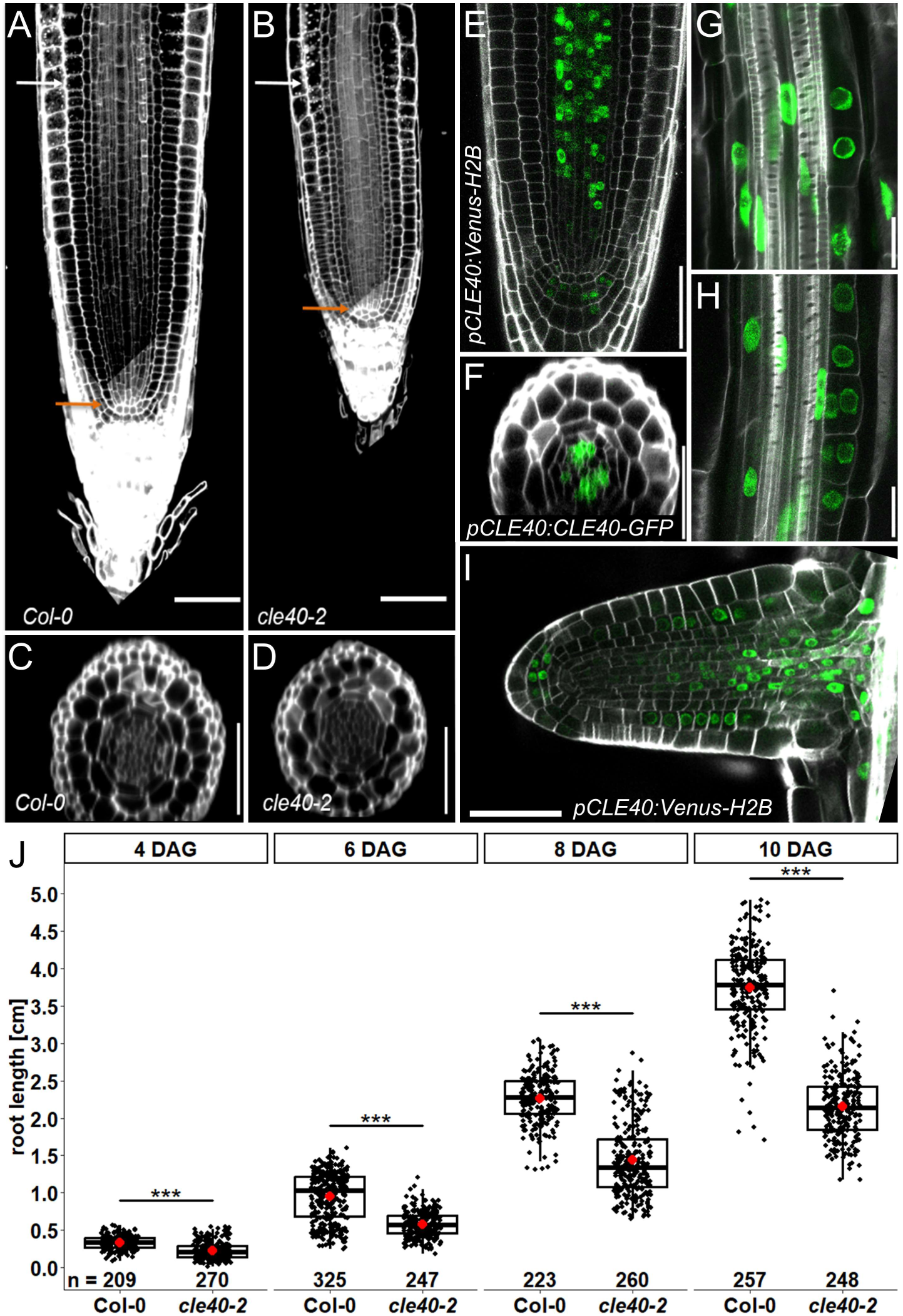
*CLE40* expression domains and *cle40-2* mutant phenotypes. **(A,B)** Representative pictures of Col-0 **(A)** and *cle40-2* **(B)** root tips at 11 DAG. **(C,D)** Representative cross sections at 150 µm proximal to the QC in Col-0 **(C)** and *cle40-2* **(D)**. **(E-I)** Expression pattern of *CLE40* reporter constructs within the procambium and the DRM **(E,F)**, lateral root primordia **(G,H)** and outgrowing lateral roots **(I)**. **(J)** Total root lengths of Col-0 and *cle40-2* at different time points (DAG). Asterisks indicate significant differences (Wilcoxon-Mann-Whitney test; signif. codes: 0 ‘***’ 0.001 ‘**’ 0.01 ‘*’ 0.05 ‘ns’). Scale bars represent 50 µm in **A-F** and **I**, and 20 µm in **G** and **H**. In **A** and **B**, orange arrows mark QC position, white arrows mark onset of TZ at the end of the PRM.

Analysis of both, the *pCLE40:CLE40-GFP* translational reporter (**60**) as well as the transcriptional reporter that targets the VENUS fluorescent protein to the nucleus (*pCLE40:VENUS-H2B*) showed *CLE40* expression in the CSCs and CCs of the DRM, and in the vasculature of the PRM starting between six to eight cells proximal to the QC (Figure 1 E, F). Optical cross sections revealed *CLE40* expression mainly in the procambium (Figure 1 F), which extended occasionally into cortex and epidermis. *CLE40* gains expression in the xylem starting precisely at the transition zone of the root (Figure S1). Furthermore, *CLE40* expression could be detected in early stages of lateral root primordia (LRP) development (Figure 1 G, H). In outgrowing lateral roots, the expression pattern resembled that of the primary root, with the exception of more cortical expression seen in early stages (Figure 1 I).

### CLE40 affects auxin signalling pathways

Elevated levels of auxin promote cell division activity resulting in a larger PRM, whereas reduced levels of auxin or a disturbed auxin distribution leads to differentiation of TA-cells, and thus to a shorter PRM (**5**,**12**,**50**). To test whether auxin signaling is affected in the PRM of *cle40* mutants, we monitored auxin dependent transcription via the synthetic auxin response reporter *DR5rev:erGFP* (**21**,**63**) and additionally analyzed *PIN1* gene expression. In the DRM, *DR5rev:erGFP* expression levels in roots of *cle40-2* mutants were not significantly different from the wild-type. However, proximal to the QC (orange box in Figure 2 A, B), where PIN1 acts to transport auxin basipetally into the root tip, *DR5rev:erGFP* activity was significantly reduced in *cle40-2* mutants (Figure 2 A, B, G). We also observed that the expression level of *PIN1* was reduced in *cle40-2* (Figure 2 C, D, H), and conclude that *CLE40* signaling promotes expression of *PIN1* and auxin signaling in the PRM. Inspecting the promoter region of *CLE40*, we found two canonical TGTCTC motifs as well as two other predicted binding sites for AUXIN RESPONSE FACTORS (ARFs) (Figure S2), which prompted us to ask whether auxin signalling or transport and *CLE40* are subject to mutual regulation. We therefore analyzed the transcriptional reporter *pCLE40:VENUS-H2B* in roots that were grown for 24 h on medium containing 1µM IAA or DMSO (mock), and found that IAA increased the fluorescent signal of the *CLE40* reporter significantly (Figure 2 E, F, I), indicating a positive feedback regulation between auxin and *CLE40*. In order to investigate the observed upregulation of *CLE40* promoter activity after auxin treatment further, we used a simplified *A. thaliana* protoplast system to analyze whether ARF5 can regulate expression at the *CLE40* promoter via a quantitative firefly luciferase assay. For this, we made use of either the full *CLE40* promoter, or a version lacking the predicted ARF5 binding region (see Figure S2). Additionally, we expressed WUS and ARF1 separately or together with ARF5 to determine whether competition or synergy occurred. We also tested if the induction with 1 µM IAA increase the transcriptional regulation by ARF5 (Figure S3). The measured values are close to the background level, which does not indicate that ARF5 is able to activate expression at the *CLE40* promoter in the tested protoplast system. Furthermore, also IAA treatment showed no clear effects, indicating that the cellular context and possibly other ARFs are responsible for the observed upregulation seen in the PRM.

**Figure 2:**
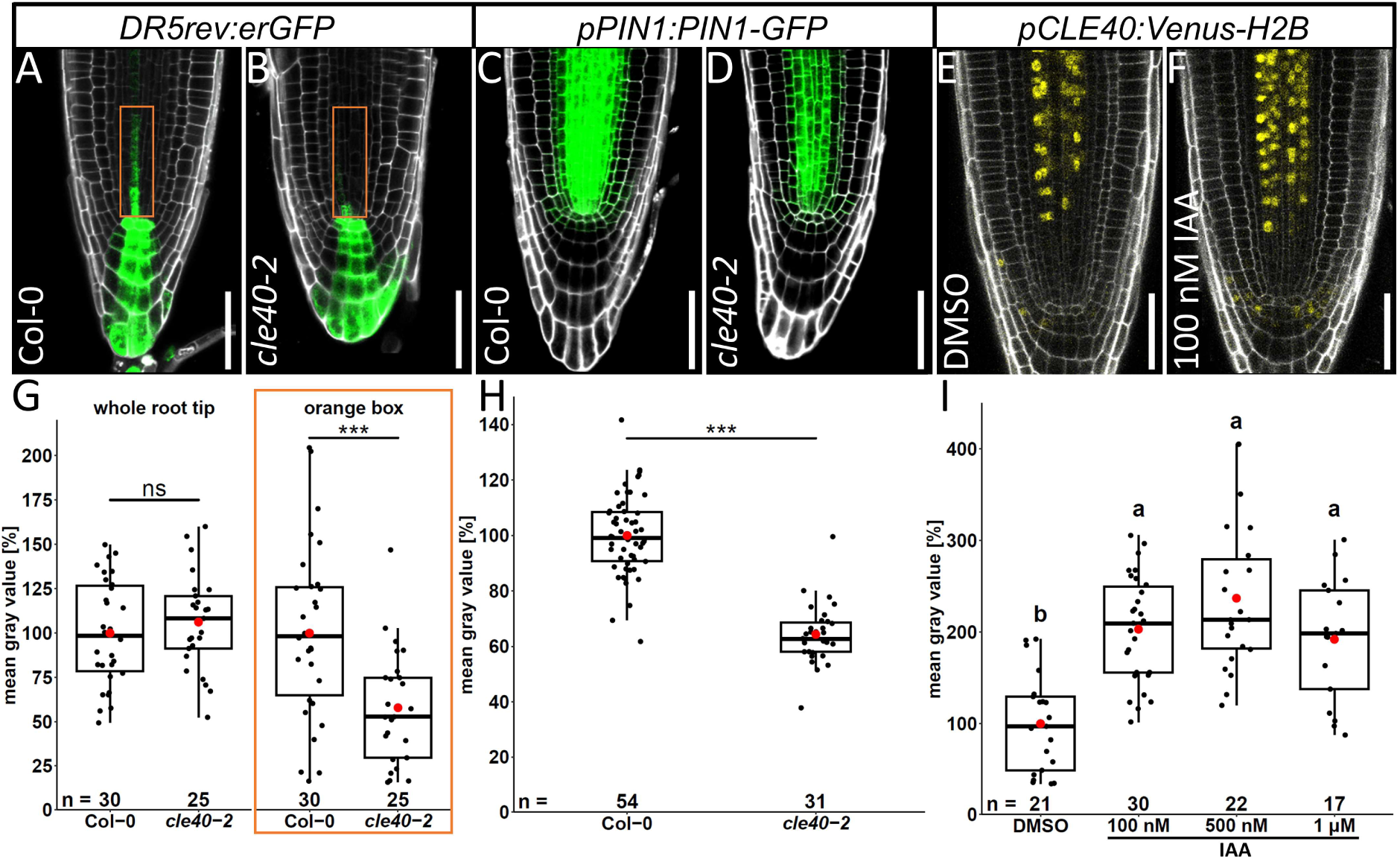
Mutual regulation between auxin and *CLE40* in the PRM. **(A,B)** *DR5rev:erGFP* expression indicating auxin maxima in Col-0 **(A)** and *cle40-2* **(B)** at 5 DAG. **(C,D)** Translational *pPIN1:PIN1-GFP* reporter activity in Col-0 **(C)** and *cle40-2* **(D)** at 5 DAG. **(E,F)** Expression of *pCLE40:Venus-H2B* after 24 h treatment DMSO (mock) or 100 nM IAA. **(G-I)** Signal quantification of reporters shown in **A-F**. In **G** and **H**, asterisks indicate significant differences (Wilcoxon-Mann-Whitney test; signif. codes: 0 ‘***’ 0.001 ‘**’ 0.01 ‘*’ 0.05 ‘ns’). In **I**, different letters indicate significant differences between groups (Kruskal-Wallis with post hoc Dunn’s test; *p*<0.05). Orange box in **G** corresponds to orange boxes in **A** and **B**. All scale bars represent 50 µm.

### CLE40 affects *PLT* gene expression levels

PLTs were previously shown to regulate mitotic activity and thereby root zonation in the PRM in a dosage dependent manner (**22**,**35**). Because multiple *plt* mutants show similar PRM defects to *cle40* mutants (**2**,**22**), we analyzed whether *PLT* expression is also altered in *cle40-2* roots. *pPLT1:erCFP* and *pPLT2:erCFP* expression was significantly downregulated in *cle40-2* compared to wild-type (Figure 3 A, B, I and E, F, J, respectively), suggesting that the root meristem defects in *cle40-2* might be caused by reduced PLT levels. To test whether the PRM defects in *cle40-2* mutants can be bypassed by increasing *PLT* expression, we assayed the effect of dexamethasone (DEX) inducible *PLTs* in *cle40-2* (**22**,**33**). *PLT1* and *PLT2* were separately overexpressed in Col-0 and *cle40-2* mutants utilizing *35S:PLT1-GR* and *35S:PLT2-GR* transgenes. After induction with DEX, the PRM size of *35S:PLT1-GR* and *35S:PLT2-GR* plants increased significantly (Figure 4, Figure S4, Table S3). Induction of *PLT1* led to a 16 % increase of PRM-size in Col-0, and a 96 % increase in *cle40-2*, while *PLT2* induction led to a 110 % increase in Col-0 and 121 % in *cle40-2*. This increase of PRM size correlated with the increase of PRM cell numbers (Table S3), indicating that expansion of the PRM resulted from accumulation of TA-cells in the meristematic region. We concluded that CLE40 signaling is required to maintain *PLT1* and *PLT2* expression levels in the root meristem, and that loss of CLE40 signaling can at least partially be compensated by increased *PLT* expression.

**Figure 3:**
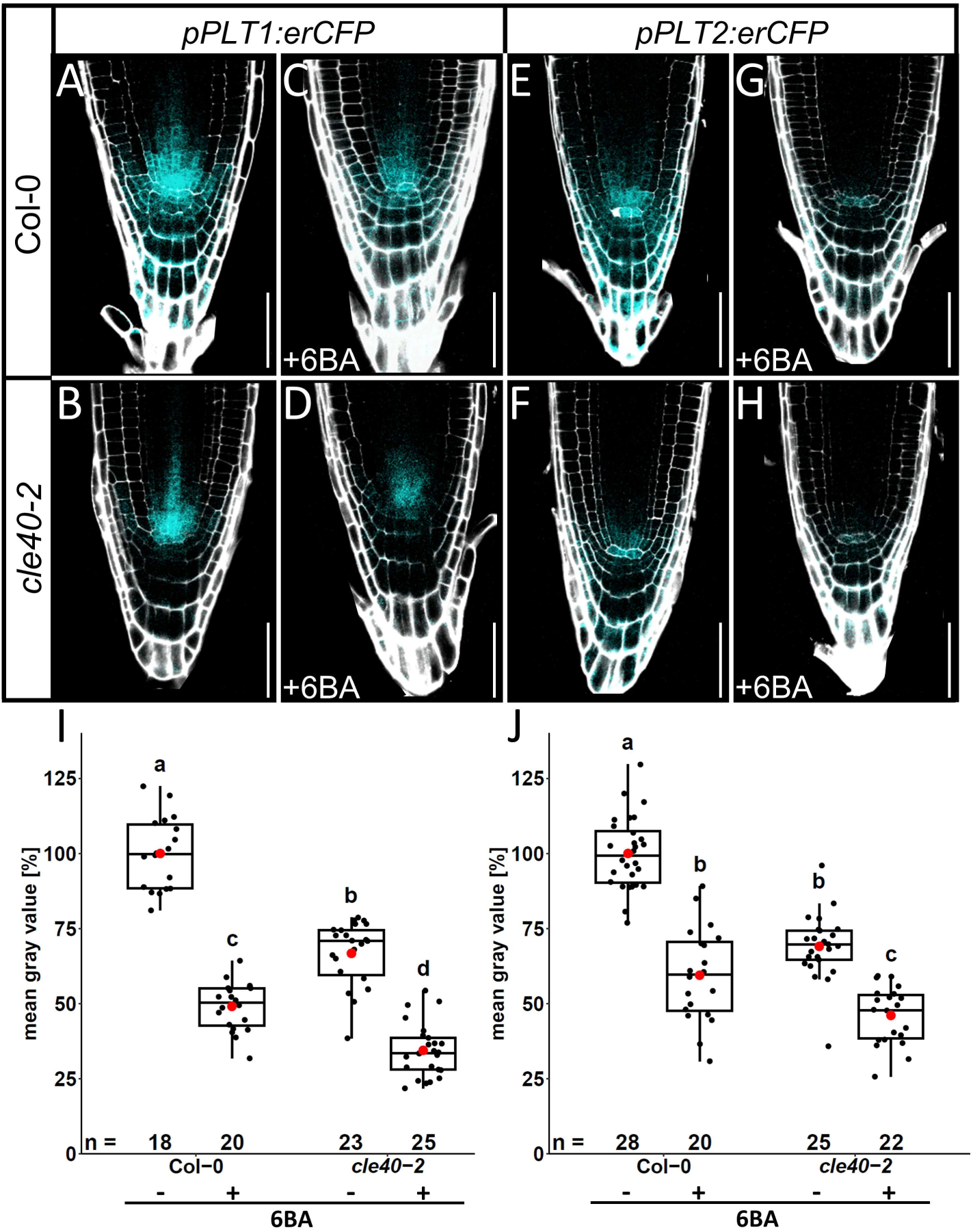
CLE40 and cytokinin affect *PLT* expression. Expression of the transcriptional reporter *pPLT1:erCFP* and *pPLT2:erCFP* in Col-0 **(A,E)** and *cle40-2* **(B,F)**, and upon treatment with 1 µM 6-BA **(C,G** and **D,H**, respectively). **(I,J)** Signal quantification of reporters shown in **A-H**. Different letters indicate significant differences between groups (Kruskal-Wallis with post hoc Dunn’s test; *p*<0.05). All scale bars represent 50 µM.

**Figure 4:**
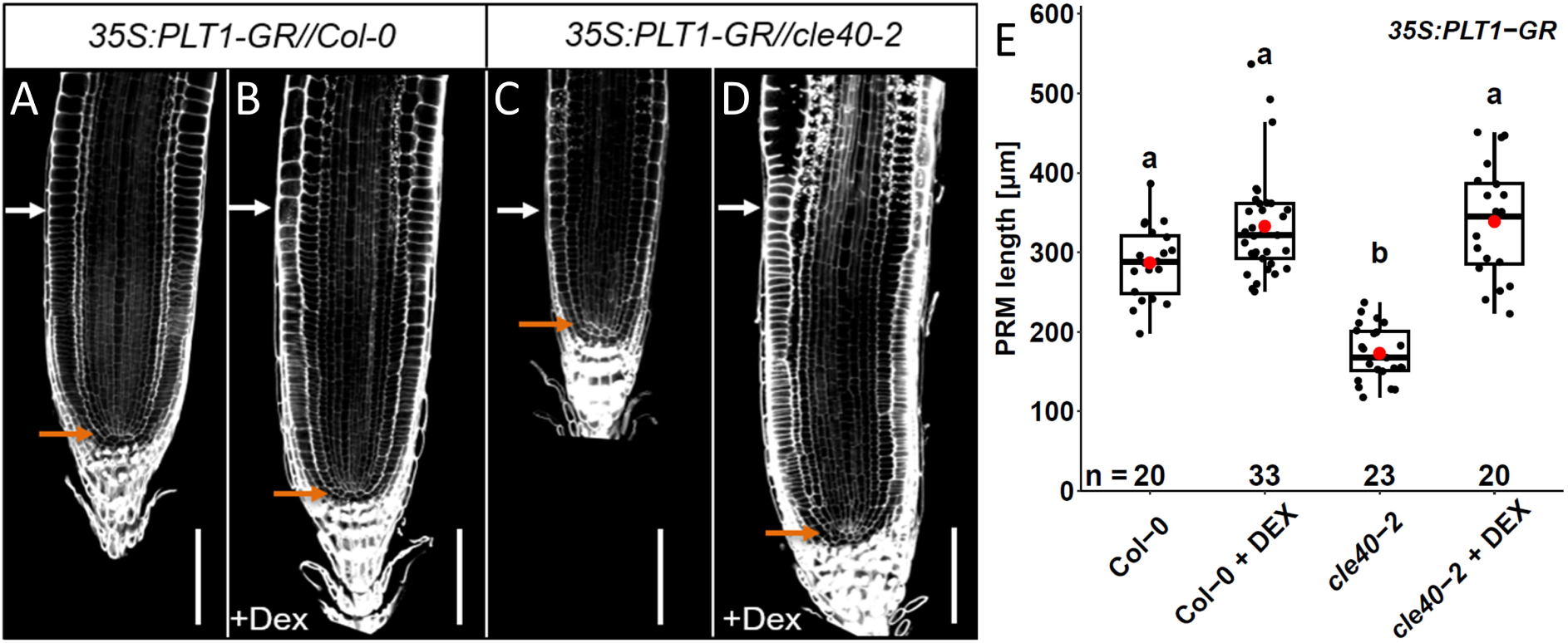
Overexpression of *PLT1* rescues PRM-defects in *cle40-2* mutants. Representative images of roots after overexpression of *PLT1* without **(A,C)** and with 3 days of 2 µM dexamethasone (DEX) treatment **(B,D)** in Col-0 and *cle40-2*, respectively. **(E)** Quantification of the PRM length dependent on overexpression of *PLT1*. Different letters indicate significant differences between groups (Kruskal-Wallis with post hoc Dunn’s test; *p*<0.05). All scale bars represent 50 µM. Orange arrows mark QC position, white arrows mark onset of TZ at the end of the PRM.

To analyze if CLE40 acts mainly through regulation of *PLT1* and *PLT2*, we studied genetic interactions between *CLE40*, *PLT1* and *PLT2*. *plt1-4,plt2-2* double mutants display a severe reduction in root length and introducing *cle40-2* led to even shorter roots, indicating additive interactions (Figure S5). We therefore concluded that CLE40 acts also upon other target genes, and not exclusively through PLT1 and PLT2.

### *CLE40* dependent control of cytokinin signaling

Cytokinin acts antagonistically to auxin on root meristem size through modulating the auxin signaling machinery (**11**,**12**,**50**). High levels of cytokinin signaling cause changes in *PIN* expression levels leading to a disturbed auxin distribution in the PRM, which results in premature differentiation of meristematic cells and therefore a short root phenotype. Therefore, *PLTs* might be indirectly regulated by cytokinin via the auxin concentration in the root tip. Hence, we asked whether CLE40 signalling outputs are mediated by cytokinin. First, we carried out a control experiment to recapitulate whether *PLT1* and *PLT2* reporters respond to increased cytokinin concentration, as shown previously by qRT-PCR (**12**). Upon cytokinin treatment with 1 µM 6-benzylaminopurin (6BA), a synthetic cytokinin, the transcriptional *PLT1* and *PLT2*-reporters were downregulated, while their general expression patterns appeared unchanged (Figure 3). The already reduced expression of the *PLT* reporter lines in the *cle40-2* background was further decreased by exogenous cytokinin addition. Next, we analyzed the expression level of *ARR5*, a negative regulator of cytokinin signaling that is rapidly upregulated in the presence of cytokinin and can therefore also serve as a readout for cytokinin signaling (**9**). In the *cle40-2* mutant background, the *pARR5:GUS* reporter showed enhanced activity (in ∼ 30 % of analyzed roots) in the PRM compared to wild-type, indicating elevated cytokinin signaling when *CLE40* is lost (Figure S6). We then asked if the cytokinin receptors CRE1 and AHK3 mediate the root length reduction of *cle40-2* mutants. While *ahk3-3* single mutants displayed root lengths similar to wild-type, *cre1-12* single and *cre1-12,ahk3-3* double mutant roots were slightly shorter (Figure 5 A). The *cre1-12,ahk3-3,cle40-2* triple mutant resembled the receptor double mutant and rescued the shorter roots of the *cle40-2* mutation, suggesting that increased signaling through cytokinin receptors is causal for the observed PRM length reduction in *cle40-2*.

**Figure 5:**
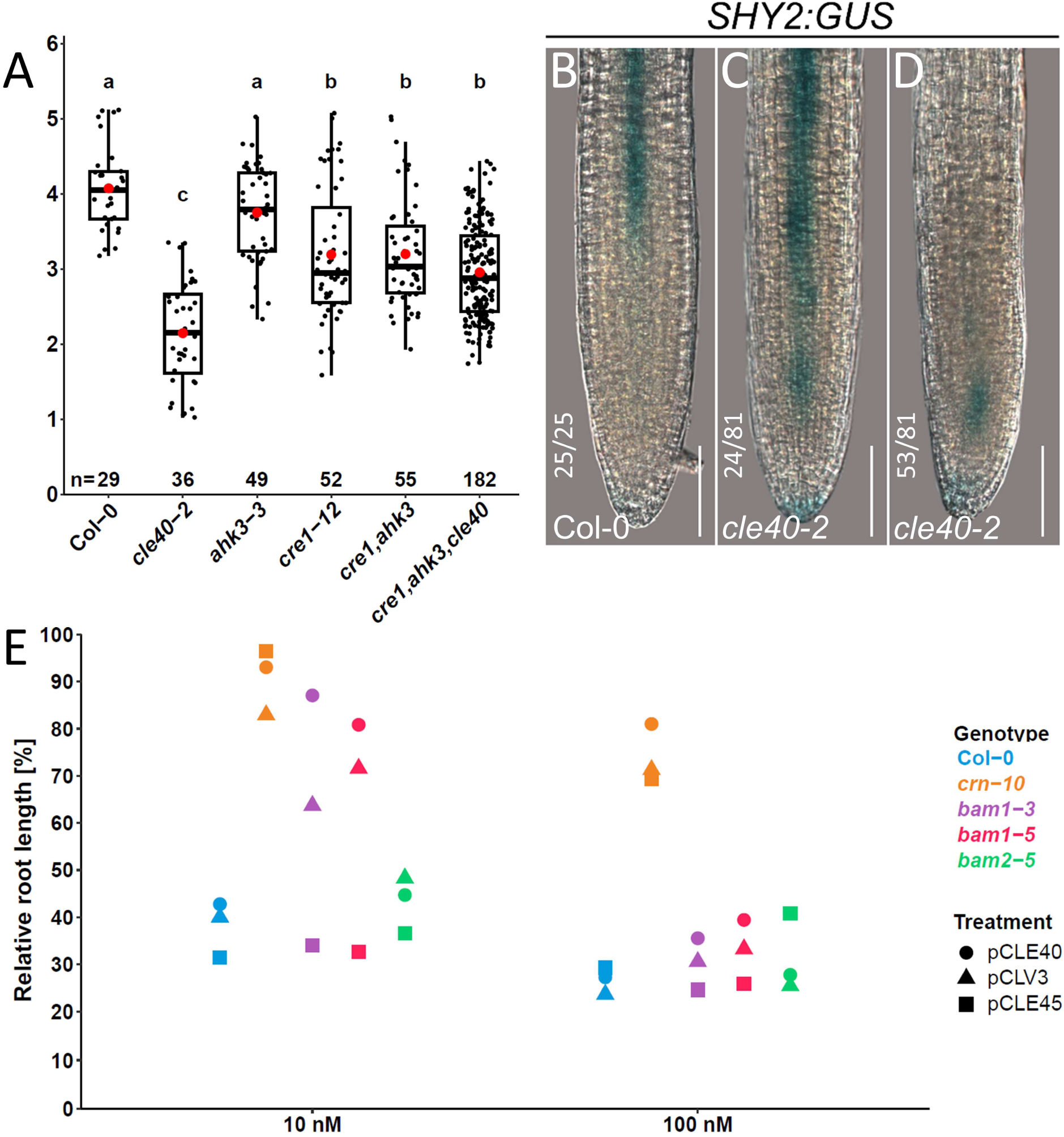
Lack of CLE40 signaling through BAM1 leads to ectopic *SHY2* activation. **(A)** Root length of indicated genotypes at 7 DAG. Different letters indicate significant differences between groups (Kruskal-Wallis with post hoc Dunn’s test; *p*<0.05). **(B-D)** In *cle40-2* mutants the *SHY2* expression domain is often found to be expanded **(C)** or shifted **(D)** towards the QC-position, compared to wild type plants **(B)** All scale bars represent 50 µM. **(E)** Relative root length at 6 DAG (*bam2-5*) or 7 DAG (all other genotypes) measured after growth on different CLE peptides compared to mock treated samples of the respective genotype. Each data point represents the average of multiple analyzed roots (n ≥ 7). Detailed measurements shown in Figure S7.

### *CLE40* signalling positions the *SHY2* expression domain

*SHY2* affects root meristem size by mediating crosstalk between cytokinin and auxin-signaling in the PRM (**12**). Loss of *SHY2* leads to more cells in the PRM and a longer root whereas *shy2* gain of function leads to shorter roots (**12**,**24**). As *SHY2* expression overlaps with *CLE40* expression at the TZ and is cytokinin inducible (**12**), it is a possible target for *CLE40* signalling. While in the wild-type *SHY2* expression starts at the TZ and is maintained at high levels in the elongation zone (Figure 5 B), in the short roots of *cle40-2* mutants, we observed that *pSHY2:GUS* expression extended to a position close to the QC (Figure 5 C), often accompanied by a reduction of *pSHY2:GUS* in the shootward regions (Figure 5 D). From these results we concluded that CLE40 signaling is required for the correct positioning and control of the *SHY2* expression domain.

### *CLE40* signaling is mediated by BAM1 in the PRM

We finally asked which receptors may transmit the CLE40 signal in the PRM to downregulate cytokinin. Previous results showed that *bam1-3* and *bam1-3,clv1-101* double mutants are partially resistant to exogenous CLE40 peptide treatments (while *clv-101* shows wild-typic sensitivity), suggesting that BAM1 is needed to perceive CLE40 (**53**). To analyze this in more detail, we grew *bam1-3*, *bam1-5* and *bam2-5* mutants on medium containing 10 or 100 nM CLE40, CLV3 or CLE45 peptide. Additionally, Col-0 and *crn-10* plants were used as controls being susceptible or highly resistant towards treatments with root active CLE peptides, respectively (**37**,**40**). As expected, root lengths of Col-0 seedlings were strongly decreased when grown on medium containing the different peptides, while *crn-10* roots showed strong resistance towards low concentrations and slightly weaker resistance to higher concentrations of all used CLE peptides (Figure 5 E and Figure S7). Both *bam1* alleles showed resistance to low levels of exogenous CLE40 and CLV3, but not CLE45 treatment, but were sensitive to high levels of all peptides. Contrarily, *bam2-5* was completely sensitive to all peptide treatments and behaved like the wildtype control. These data suggest that BAM1 is specifically binding CLE40 - and possibly other, but not all CLE peptides - in the PRM; at very high and possibly unphysiological concentrations of CLE40, other receptors might mediate signalling.

## Discussion

*cle40* mutants were described to develop shorter roots than the wild-type (**28**,**43**). Initially this decreased root length was assumed to be the result of enhanced root waving and later studies suggested that CLE40 regulates cell differentiation in the PRM (**43**). To elucidate the specific role of CLE40, we analyzed total root length, growth speed and the PRM structure of *cle40* loss of function mutants in more detail. We found that the reduction of total root length was caused by premature differentiation of PRM cells, which leads to an overall slower growth and growth termination ahead of the wild-type. The size of cortical PRM cells was unchanged in comparison to Col-0, thus a reduced root length due to smaller cells could be excluded (Table S1). In addition to a reduced number of cells along the longitudinal axis, we counted less epidermal, and vascular cells in root cross sections, thus explaining the smaller perimeter in *cle40-2* roots (Table S2). Introducing a copy of the *CLE40* gene under its native promoter could restore wild-type root length, demonstrating that *CLE40* is required for normal root growth (**53**). Interestingly, when looking at the expression domain in root cross sections, we found that *CLE40* promoter activity is restricted to procambial cells in the PRM, but gains additional activity in xylem cell files in the TZ (Figure S1). Presumably, the CLE40 promoter contains unknown cis-regulatory elements that could either serve to repress its activity in developing xylem cells prior to entering the transition zone or specifically activate expression once the cells start differentiating.

The *CLE40* expression pattern (Figure 1) suggests that it regulates PRM size from the stele. Previously, we showed that *CLE40* exerts a stem cell inhibiting function in the DRM by restricting CSC fate via the RLKs ACR4 and CLV1 (**59**,**60**). The finding that *CLE40* exhibits the opposite function in the PRM could be explained by differing receptor availabilities between these domains. *ACR4* is most strongly expressed in tissues of the DRM, beneath the QC. In the PRM it is only weakly expressed in epidermal cell layers and in pericycle cells during lateral root primordia initiation (**10**,**23**). Similarly, *CLV1* expression was found in the DRM (**59**) as well as phloem companion cells (**3**). Therefore, both *ACR4* and *CLV1* expression do not strongly overlap with *CLE40* in the PRM, indicating that other receptors that are expressed in the PRM might perceive the CLE40 signal there. It was shown that CRN and CLV2 can be targeted by CLE40, but only in CLE40 overdose conditions, such as high-level ectopic application (**19**,**28**,**40**,**43**,**47**). Other putative receptors for CLE40 in the PRM are the BARELY ANY MERISTEM (BAM) receptors, which are closely related to CLV1 (**14**). The published transcriptional reporters of *BAM1* differ in their expression pattern in the root tip, but are both expressed within the stele (**18**,**42**). Therefore, *BAM1* expression extensively overlaps with that of *CLE40* in the PRM. Moreover, *bam1-3* (but not *clv1-101*) roots tend to be slightly shorter than wild-type and were shown before to be partially resistant to CLE40 peptide treatment (**53**). By analyzing root length reduction in different genetic backgrounds and with different CLE peptides we found that two *bam1* mutants are specifically insensitive towards treatment with low concentrations (10 nM) of CLE40 or CLV3, but sensitive to higher concentration (100 nM) of these peptides. Additionally, both alleles were fully sensitive to the root active peptide CLE45 (Figure 5 E and Figure S7). The peptide sequences of CLE40 and CLV3 are more similar compared to the one of CLE45. Especially in the N-terminal part of the peptide, which was suggested to be the main recognition and docking point between BAM1 and CLE9 before (**48**), CLE45 comprises more basic amino acid residues which may render it unable to bind to BAM1. We further found that a mutation in the closely related receptor BAM2 did not lead to any resistance towards any of the peptides. We therefore conclude that BAM1, under normal conditions, perceives CLE40 in the PRM.

It was shown that reduced auxin levels can lead to a shortened PRM (**5**,**12**) and that an interplay of auxin, PLTs and cytokinin response regulators are needed for the correct positioning and stabilization of the TZ (**5**,**15**,**51**). To link these observations to the shortened PRM of *cle40-2* mutants, we asked whether CLE40 regulates components of the auxin or cytokinin signaling machineries. Previously, a transcriptome analysis of *cle40* roots identified several auxin signaling related genes which showed differential expression compared to the wild-type. However, an explicit direction of regulation could not be determined by this method as genes with similar function showed opposing regulation and it was therefore assumed that CLE40 differentially affects auxin signaling (**43**).

Our analysis of *DR5rev:erGFP* expression levels revealed a reduction of auxin signaling in the PRM when CLE40 is missing (Figure 2 A, B, G). Furthermore, *PIN1* as well as *PLT1* and *PLT2* levels were reduced compared to wild-type (Figure 2 C, D, H, Figure 3). Since it was reported that PLTs are needed for *PIN* transcription and expression levels of multiple *PIN*s are strongly reduced in higher order *plt* mutants (**5**,**22**), it is likely that not only *PIN1* but also other *PINs* exhibit a reduced expression in *cle40-2* mutants, although not analyzed here. Several single and high-order mutants of *PIN* genes generate shorter roots (**5**), indicating that partial expression changes might be sufficient to lead to similar defects in root development. The reduced levels of *PLT1* and *PLT2* in *cle40* mutants additionally suggest that CLE40 also affects cell division activity of TA-cells through modulating *PLT* levels, which were shown to have an inhibiting effect on differentiation (**35**). The fact that overexpression of *PLT* genes rescued the PRM defects of *cle40* roots (Figure 4, Figure S4, Table S3) supported our hypothesis that *CLE40* affects the *PLT* gene regulatory network. However, the root length of *plt1-4,plt2-2* double mutants was further reduced in *plt1-4,plt2-2,cle40-2* triple mutants (Figure S5), suggesting that *CLE40* promotes PRM size partially independent of *PLT*s.

Cytokinin acts upstream of auxin signaling in the root and our transcriptome analysis suggested an inhibiting role of CLE40 on cytokinin signaling (**43**). The downregulation of *PLT* expression after cytokinin (6BA) treatment, which showed an additive effect with the already reduced levels in the *cle40-2* background, supported this hypothesis (Figure 3). In line with this, *ARR5* expression, used as readout of cytokinin activity, was upregulated in *cle40-2* roots (Figure S6).

Introducing *cle40-2* into the cytokinin receptor double mutant *cre1-12,ahk3-3* resulted in the same root length observed in the double mutant alone (Figure 5 A), which indicated that the effects of *CLE40* signaling are transmitted via these receptors. Furthermore, the changed expression pattern of *SHY2*, which extended or shifted to a position close to the QC (Figure 5 B-D), indicated that the modified cytokinin signaling can lead to a reshaping of auxin distribution and a shift in TZ positioning.

## Conclusion

We summarized our findings in a model (Figure 6). Altogether, our experimental data suggest that CLE40, under normal conditions, downregulates cytokinin signaling in the PRM. This happens via a yet unresolved mechanism comprising the perception of CLE40 by the LRR-RLK BAM1 in the stele of the PRM. In *cle40-2* mutants, this control is missing, leading to higher cytokinin signaling through AHKs and an ectopic activation of *SHY2* expression. SHY2 in turn reshapes the auxin distribution in the root, which ultimately leads to a relocation of the TZ to a position closer to the QC.

**Figure 6:**
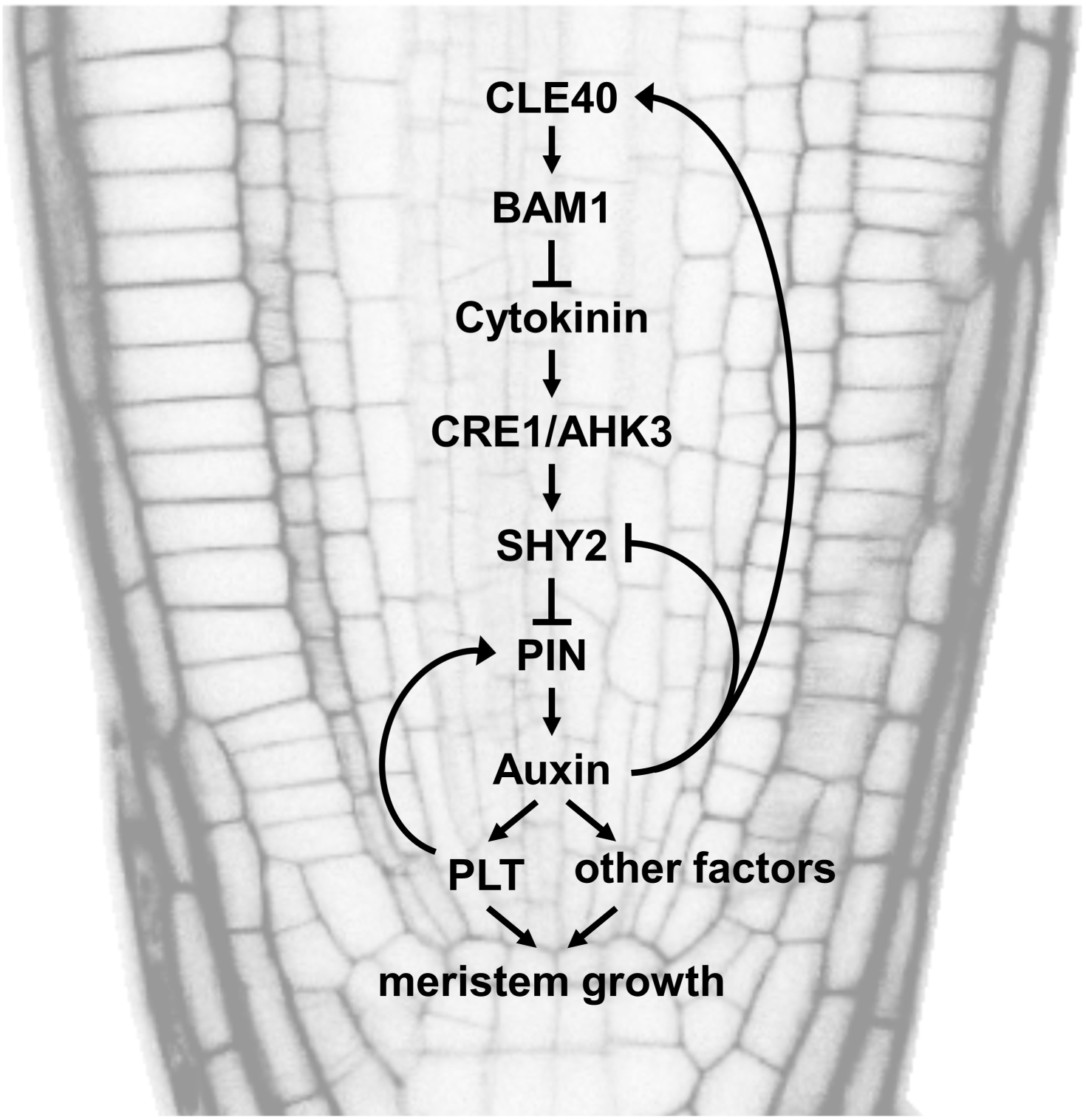
Simplified model for *CLE40*-dependent regulation of the PRM. Arrow-headed lines indicate activation and bar-headed lines indicate inhibition.

The establishment of root meristem and stabilization of the TZ has been studied extensively (**57**,**62**). With our work, we add a new layer to this regulation based on a mobile peptide, which itself is modulated by auxin. Future work will be needed to elucidate which subsequent mechanisms are activated after the perception of CLE40 by BAM1 that lead to lower cytokinin signaling activity. Furthermore, CLE40 signaling affecting the positioning of the TZ, but itself being sensitive to signals in this region - shown by the additional expression gained in the xylem - adds another level of complexity to this system.

## Materials & Methods

### Plant Growth Conditions

All plants were grown in climate chambers under controlled conditions at 21 °C. These chambers were set on continuous light. Seedlings used for root growth analysis or hormone and peptide assays were surface sterilized using chlorine gas, soaked in 0,1% (w/v) agarose and incubated in darkness for two days on 4 °C before grown on vertically oriented 1/2MS-plates (0.5x Murashige and Skoog (MS) medium with Gamborgs No. 5 vitamins, 0.5 g/L 2-(N-morpholino) ethanesulfonic acid (MES), 1 % (w/v) sucrose and 1.2 % (w/v) plant agar, containing desired supplements (as indicated in figures/figure descriptions). For peptide treatments, peptides synthesized by Davids Biotechnologie GmbH (Regensburg, Germany) were used. CLV3: RTVHypSGHypDLPHHH; CLE40: RQVHypTGSDPLHHK; CLE45: RRVRRGSDPIHN.

### Plant Materials

*Arabidopsis thaliana* ecotypes Columbia-0 (Col-0) and Wassilewskija (WS) were used. *cle40-2* mutants (Col-0) were identified as described in (**60**). *plt1-4,plt2-2* (WS) were obtained from the Nottingham Arabidopsis Stock Centre (NASC, UK) and were confirmed by PCR-based genotyping. *cre1-12*, *ahk3-3* as well as the receptor double mutant were kindly provided by Eva Benkova (**27**). The *pCLE40:Venus-H2B* and *pCLE40:CLE40-GFP* (Col-0) lines were previously described in (**53**,**60**), respectively. *35S:PLT2-GR* (Col-0) was kindly provided by Ben Scheres. *pSHY2:GUS* (Col-0) was kindly provided by Sabrina Sabatini. *pPIN1:PIN1-GFP* (Col-0) and *DR5rev:erGFP* (Col-0) were kindly provided by Jiri Friml. *pPLT1:erCFP* (Col-0) and *pPLT2:erCFP* (Col-0) were obtained from (**22**). The coding sequence of *PLT1* (except stop codon) was amplified from genomic DNA (*A. thaliana*, Col-0). The sequence of the glucocorticoid receptor (GR) was amplified from transgenic plants carrying a *35S:PLT2-GR* transgene. Both fragments were separately inserted in the pENTR-D-TOPO entry vector via directed TOPO-cloning and fully sequenced. *PLT1* was then amplified from the pENTR-D-TOPO-backbone including the 5’ located attL1 site and the GR sequence was amplified from the pENTR-D-TOPO-backbone, including the 3’ located attL2 site. Both fragments were finally fused with an overlap extension PCR. The result was a fused *PLT1-GR* sequence, flanked by Gateway compatible attL1 and attL2 sites. The PCR-fragment was then inserted in the destination vector pMDC32 by a Gateway LR reaction. The generated expression vector was verified via sequencing and named *35S:PLT1-GR*. All used primers and their sequences are listed in Table S4.

### In planta transformation of T-DNA into the genome of A. thaliana plants

Transformation of *A. thaliana* plants with transfer-DNA (T-DNA) was carried out using the floral dip method as described in (**6**).

### Expression Analyses

Beta-Glucuronidase (GUS) activity in root tissues was detected as described in (**60**). The stained roots were incubated overnight in clearing solution containing 70% (w/v) chloral hydrate and 10% (v/v) glycerol.

### Microscopy

Nomarski (DIC) microscopy was carried out using an Axioskop 2 mot plus microscope with Nomarski-optics (Carl Zeiss, Jena). An attached Axiocam (Carl Zeiss, Jena) was used to record digital pictures using the corresponding Axio Vision software (AxioVs40). Confocal imaging was carried out using Zeiss LSM 510 Meta, LSM 780, or LSM 880 laser scanning microscopes (CAi, HHU). Counterstaining of cell walls was achieved by mounting seedling roots in 10 μM propidium iodide or by staining overnight with SCRI Renaissance 2200 (SR2200) (**41**) after PFA fixation and optical clearing using ClearSee (**34**). SR2200 was excited at 405 nm and emission detected at 410-460 nm. GFP was excited at 488 nm and emission detected at 490-560 nm. Venus was excited at 514 nm and emission detected at 520-560 nm. CFP was excited at 458 nm and emission detected at 490-530 nm. Propidium iodide was excited at 561 nm and emission detected with a long pass at 575nm.

### Root length measures and cell counting

Root length analysis was performed and optical cross sections were generated using ImageJ/Fiji (**52**,**54**). PRM length was defined as the distance from the QC to the first elongated cortex cell. Within root cross sections, cell files were manually counted.

### Luciferase assay

Protoplasts were isolated from ∼14 days old A. thaliana plants sterile grown on square SCA medium plates. The seedlings were grown in long day conditions (16 h light, 8 h darkness) at 22 °C. The plant material was cut with a scalpel upwards of the root and transferred to a round petri dish filled with 10 mL MMC buffer. The plants were cut into small strips and incubated overnight in darkness. The plant material was digested with an enzyme mix containing 0.5 % cellulose Onozuka R10 and macerozyme R10 added to the MMC medium. After ∼18 h the lysate was carefully mixed and transferred into a 50 mL tube, passing through a 70 µm pore sieve and subsequently centrifuged at 100 rcf. After centrifugation the enzyme containing medium was removed and the protoplast pellet was resuspended in MSC buffer. After transferring 10 mL into a round bottom tube the solution was carefully overlaid with MMM solution to create two layers. Following a short centrifugation at 80 rcf the protoplasts located at the interphase of the two buffers were collected in a separate round bottom tube filled half with W5 buffer. After all protoplasts were collected, the total amount was determined via counting inside a counting chamber. For the transformation, the protoplasts were diluted to ∼500.000 per 100 mL in MMM buffer. In non-treated 6 well plates the plasmid DNA (total of 30 µg) for the transformation was collected and mixed in a total volume of 20 µL MMM for each well. In each well the 100 µL protoplasts were added to the DNA and mixed alongside the rim of the well. After a short wait, 120 µL polyethylene-glycol (PEG) was added dropwise to the DNA-protoplast mix. After 8 min, 120 µL MMM and 2×720 µL W5 with 2% FBS and 0.05% Ampicillin were added and the mixture well mixed to dilute the PEG. After transformation the protoplasts were kept in darkness for at least 20 h or induced with 1 µM IAA after 4 h and incubated 18 h further.

To measure luciferase activity, the protoplasts were transferred from the 6 well plate into a 96 well plate. Each well/transformation had four technical replicates. Each well contained 80 µl protoplast solution and 20 µL luciferase substrate added directly before measuring. The measurement was performed at TriStar multi-platereader with 29 repeats over 20 min time period. Further methodological details, as well as all buffer and luciferase substrate compositions, can be found in (**36**).

### Statistical testing and graphs

Data were tested for normal distribution (Shapiro-Wilk test and visual data inspection) and homogeneity of variance (Levene’s test). Subsequently, data was analyzed with either parametric or non-parametric testing methods, indicated in the figure descriptions. All statistical analyses were performed and all graphs were generated using R Statistical Software (**46**) in the RStudio environment (**45**). Furthermore, the packages ggplot2 (v3.5.1, (**65**)), ggsignif (v0.6.4, (**1**)), car (v3.1.3, (**20**)), coin (v1.4.3, (**29**)) and PMCMRplus (v1.9.12, (**44**)) were used.

### Accession Numbers

Sequence data from this article can be found in the Arabidopsis Genome Initiative or GenBank/EMBL databases under the following accession numbers: *AHK3* (AT1G27320), *BAM1* (AT5G65700), *BAM2* (AT3G49670), *CLE40* (AT5G12990), *CRE1* (AT2G01830), *CRN* (AT5G13290), *PIN1* (AT1G73590), *PLT1* (AT3G20840), *PLT2* (AT1G51190), *SHY2* (AT1G04240).

## Acknowledgements

We would like to acknowledge the Center for Advanced Imaging (CAi) at Heinrich Heine University for providing access to the Zeiss LSM 780 and Zeiss LSM 880 microscope systems (DFG-INST 208/551-1 FUGG and DFG-INST 208/746-1 FUGG). R.S. and M.Z. received funding from the DFG through iGRAD-PLANT (GRK 1525) and CEPLAS (EXC 2048).

## Statistical testing and graphs

**Figure S1:**
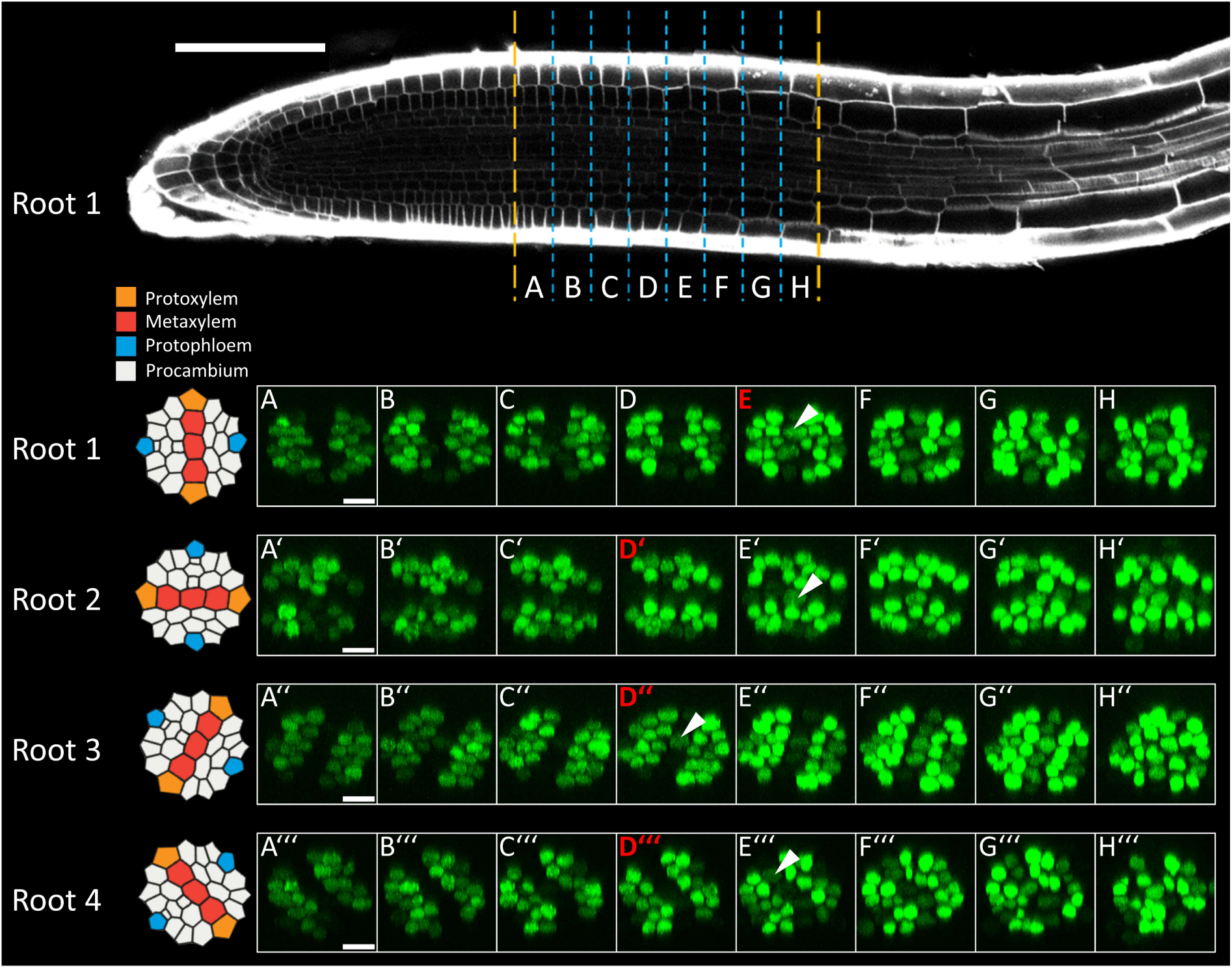
Change of *CLE40* expression pattern in developmental gradient. Maximum intensity projections of optical cross sections showing the change of *CLE40* expression in four separate roots. Each projection covers a range of 22-26 µm on the longitudinal axis. Onset of the transition zone is highlighted by a red figure label. *CLE40* is expressed in procambial cells in the PRM. Within roots 1 and 3, the onset of the transition zone directly correlates with the additional activation of *CLE40* in xylem cells (marked by white arrow heads), while in roots 2 and 4, the activation can be observed in the following section. Scale bars represent 100 µm (root tip) and 10 µm (optical cross sections). Schematic representations of root cross sections were taken and adapted from (**16**).

**Figure S2:**
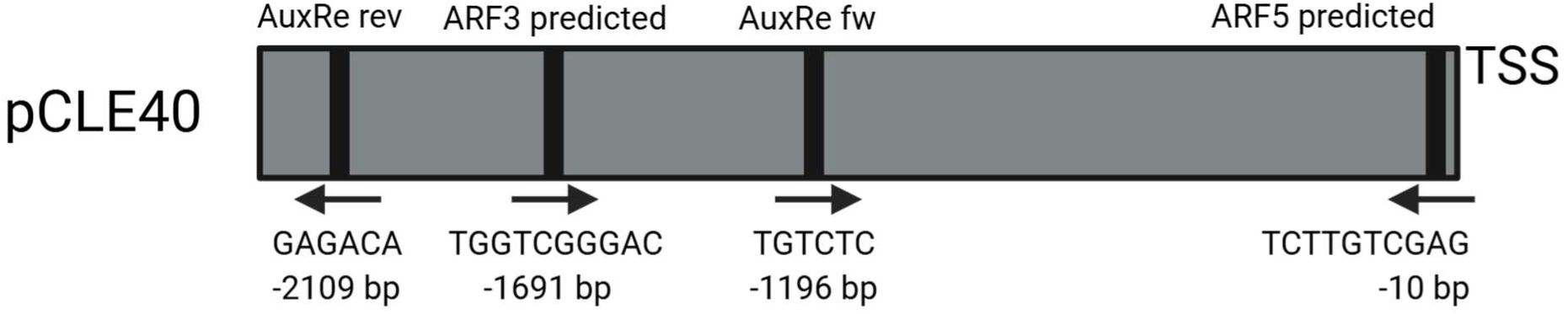
*CLE40* promoter scheme. Potential auxin responsive cis-elements in the promoter of *CLE40*. ARF binding sites were predicted using PlantTFDB. The two AuxRe elements were identified by searching for the TGTCTC motif.

**Figure S3:**
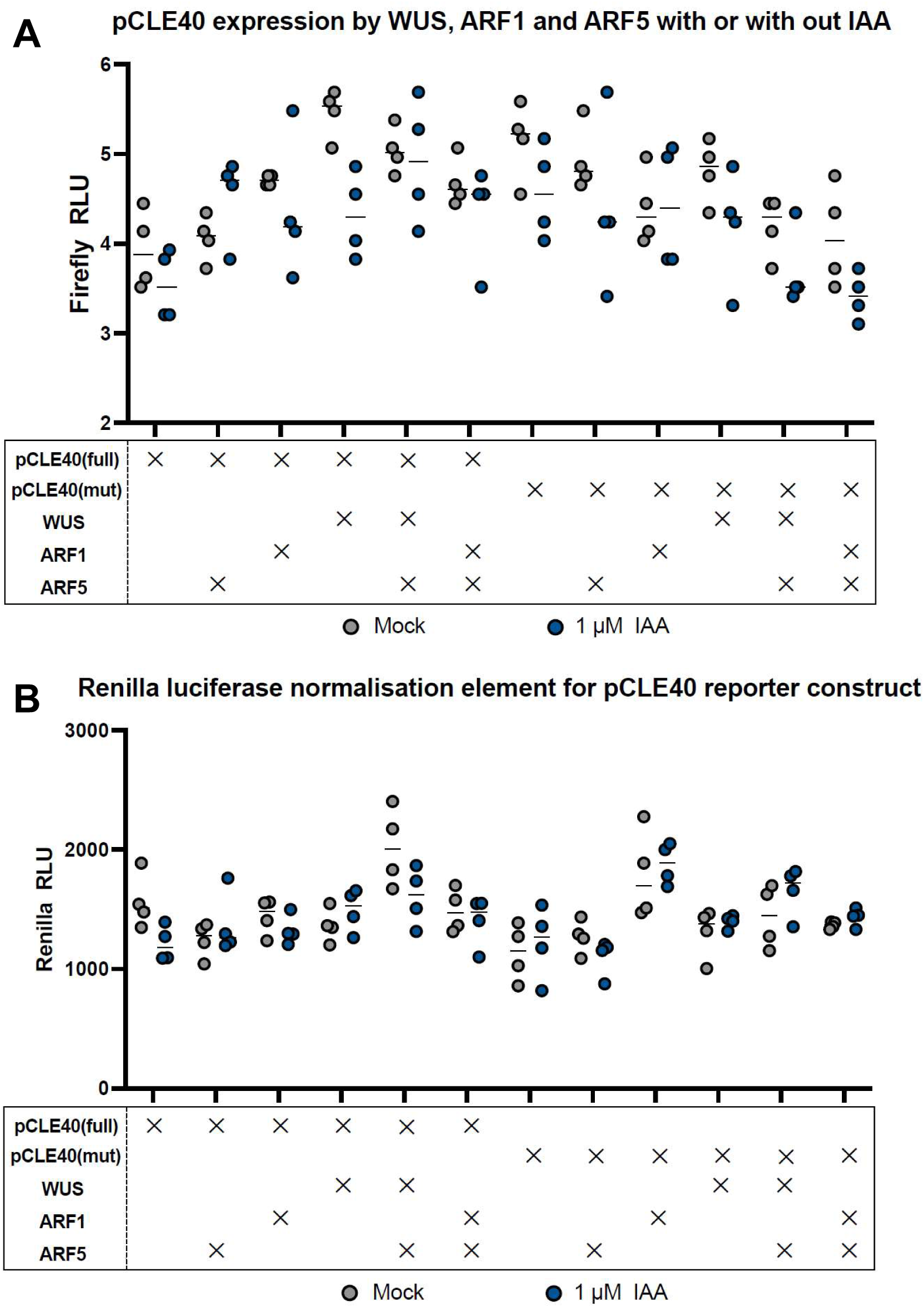
Mode of function ARF5 and WUS activation experiment in *A. thaliana* protoplasts. **(A)** The transcription factors ARF5, ARF1 and WUS were tested for their ability to initiate transcription at the tested binding site. The reporter construct consists of the Firefly luciferase reporter gene under the control of the *CLE40* promoter or the *CLE40* promoter lacking the predicted ARF5 binding site. The reporter is co-expressed with plasmids constitutively expressing WUS, ARF1 or ARF5, or in combination, via the strong 35S promoter. 4 hours after transfection, the protoplasts were split into two wells and treated either with dH2O (mock) or with 1 µM IAA. 18 hours after induction, 4 technical replicates per setup were measured for luminescence using a plate-reader. **(B)** Additionally, constitutively expressed Renilla luciferase was measured as normalization.

**Figure S4:**
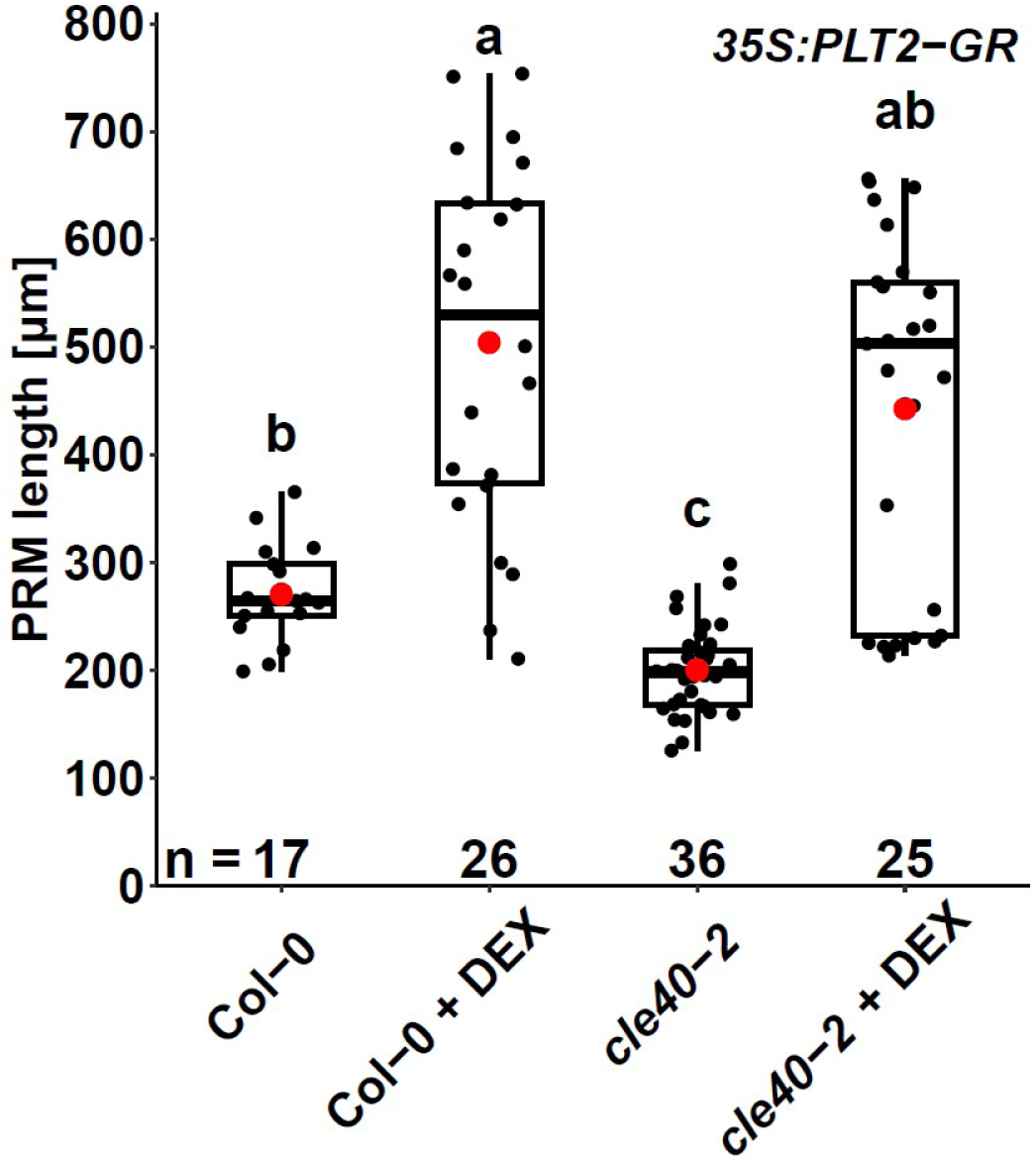
Overexpression of *PLT2* rescues PRM-defects in *cle40-2* mutants. Quantification of the PRM length dependent on overexpression of *PLT2* with or without 3 days of 2 µM dexamethasone (DEX) treatment. Different letters indicate significant differences between groups (Kruskal-Wallis with post hoc Dunn’s test; *p*<0.05).

**Figure S5:**
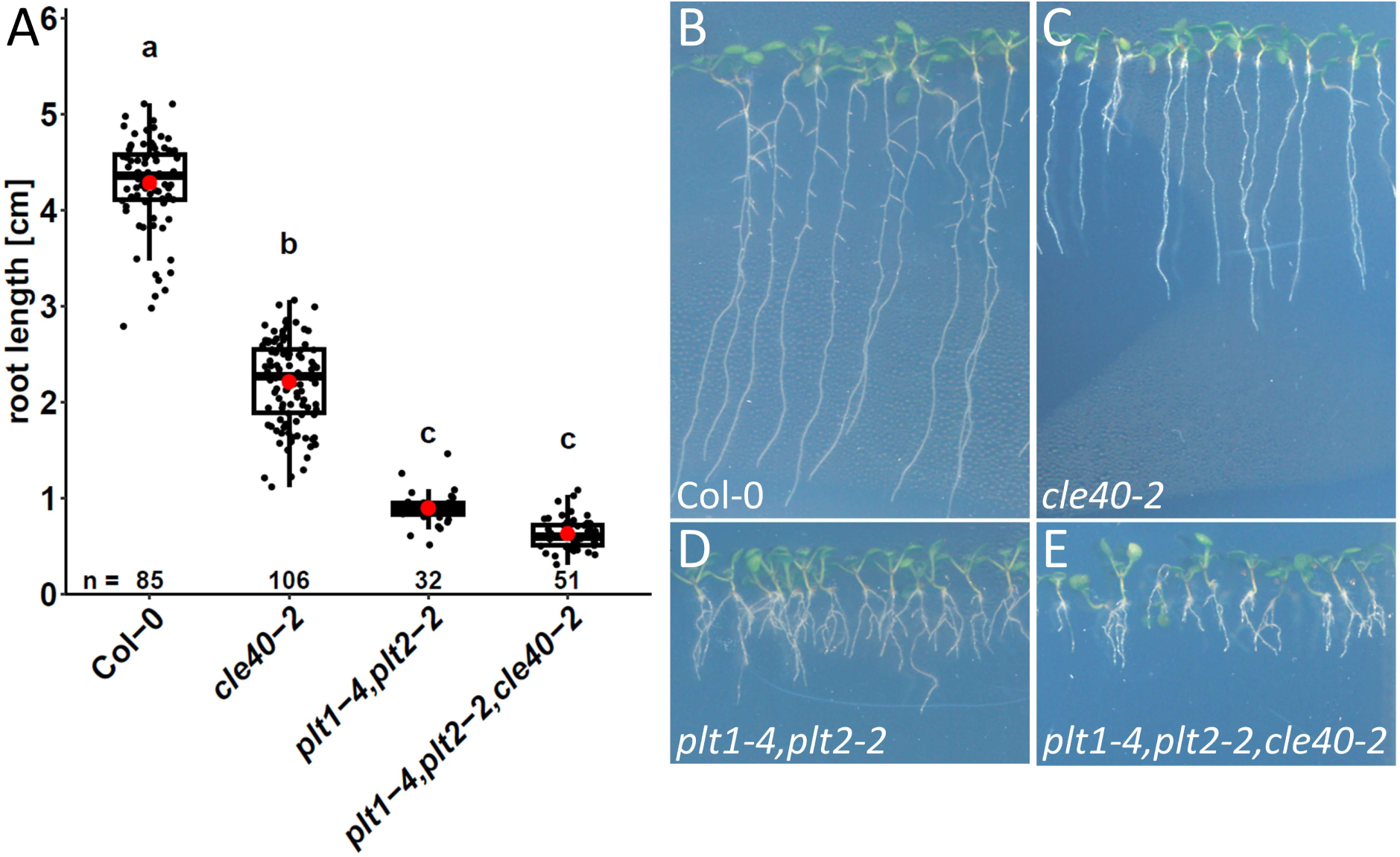
*CLE40* does not exclusively act through *PLTs*. **(A)** Root lengths of wild-type and mutant plants at 10 DAG. Different letters indicate significant differences between groups (Kruskal-Wallis with post hoc Dunn’s test; *p*<0.05). *p*-value for the comparison between *plt1-4,plt2-2* double mutant and *plt1-4,plt2-2,cle40-2* triple mutant is approximately 0.07, as the small difference (0.90±0.17 cm and 0.63±0.16 cm) is masked by larger differences to the other genoytpes. A specific pairwise analysis between these two genotypes (Wilcoxon-Mann-Whitney test) reveals a significant difference (*p*=1.472e-10). **(B-E)** Representative roots of measured genotypes.

**Figure S6:**
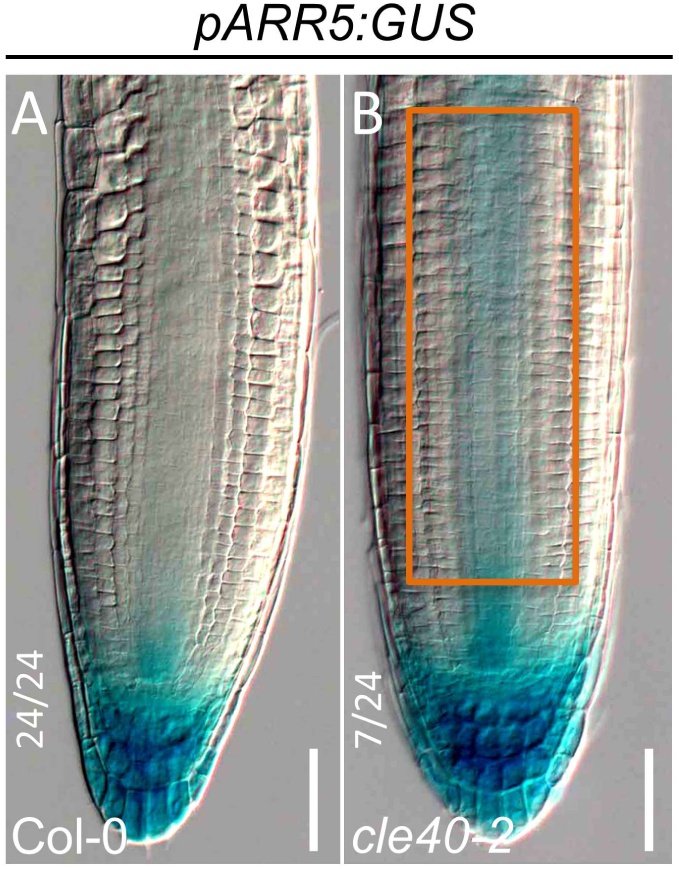
*ARR5* is upregulated in *cle40-2* roots. Representative images of *pARR5:GUS* stainings in Col-0 **(A)** and *cle40-2* **(B)** roots. Increased vascular signal in *cle40-2* was observed in 7 out of 24 analyzed roots.

**Figure S7:**
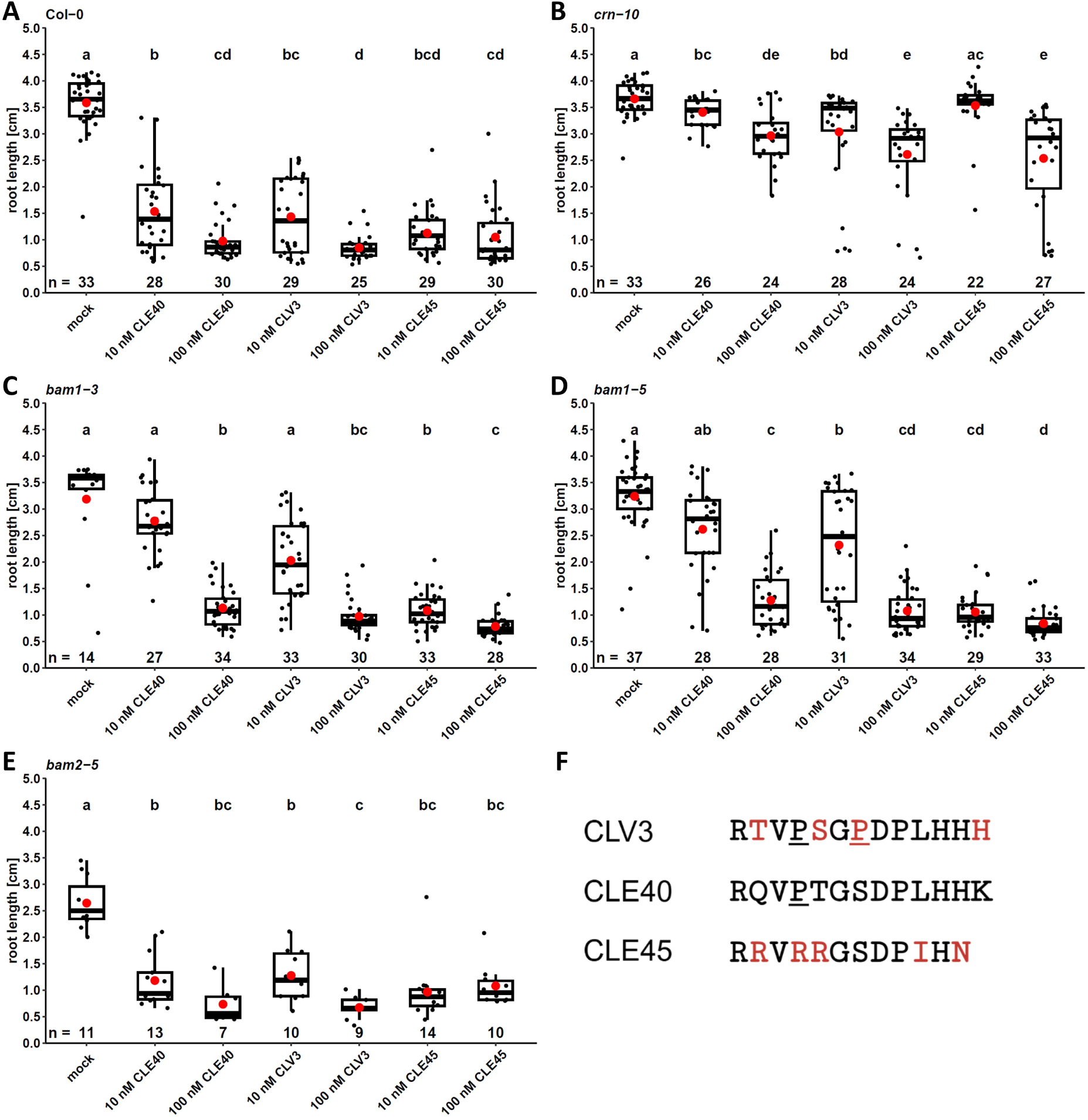
*bam1* alleles are insensitive to low concentrations of pCLE40 and pCLV3. **(A-E)** Root length of seedlings grown on mock or CLE peptide containing medium at 6 DAG (*bam2-5*) or 7 DAG (all other genotypes). Different letters indicate significant differences between groups (Kruskal-Wallis with post hoc Dunn’s test; *p*<0.05). **(F)** Sequence comparison of the used CLE peptides. Amino acid differences of CLV3 or CLE45 in comparison to CLE40 are highlighted in red. Underscored <u>P</u> refers to Hydroxyproline.

**Table S1:** Analysis of PRM length and cortex cell measures in longitudinal sections at 6 DAG old Col-0 (n=31) & *cle40-2* (n=46) plants. *p*-values were calculated using Student’s *t*-tests.

|  | **Col-0** | ***cle40-2*** | ***p*-value** |
| --- | --- | --- | --- |
| PRM length [$\mu\text{m}$] | 285.73 ± 40.61 | 186.03 ± 31.70 | 2.7e-19 |
| cortex cell number | 41.26 ± 4.94 | 26.04 ± 4.52 | 3.1e-22 |
| mean cortex cell length [$\mu\text{m}$] | 6.95 ± 0.76 | 7.18 ± 0.69 | 0.17 |

**Table S2:** Analysis of root perimeter and cell file numbers observed in PRM cross sections at 6 DAG old Col-0 (n=21) and *cle40-2* (n=20) plants. *p*-values were calculated using Wilcoxon-Mann-Whitney tests.

|  | **Col-0** | ***cle40-2*** | ***p*-value** |
| --- | --- | --- | --- |
| perimeter [$\mu\text{m}$] | 346.26 ± 18.03 | 301.55 ± 22.94 | 4.7e-99 |
| epidermal cells | 22.76 ± 1.57 | 18.25 ± 2.02 | 2.7e-8 |
| cortical cells | 8.10 ± 0.29 | 8.25 ± 0.43 | 0.2379 |
| endodermal cells | 8.76 ± 0.61 | 8.30 ± 0.46 | 0.01986 |
| vascular cells | 45.24 ± 2.88 | 39.42 ± 3.12 | 9.8e-7 |

**Table S3:** Comparison of PRM length and cell numbers in mock or DEX treated plants expressing *PLT1-GR* or *PLT2-GR*. Pairwise comparison *p*-values were calculated using Wilcoxon-Mann-Whitney tests.

| genotype | T-DNA | treatment | PRM length<br>[μm] | standard deviation<br>[μm] | *p*-value | n |
| --- | --- | --- | --- | --- | --- | --- |
| Col-0 |  | - | 302.99 | 39.23 | 0.35 | 12 |
| Col-0 |  | DEX | 324.80 | 50.13 |  | 18 |
| *cle40-2* |  | - | 192.97 | 28.12 | 0.06 | 27 |
| *cle40-2* |  | DEX | 212.30 | 34.30 |  | 19 |
| Col-0 | *35S:PLT1-GR* | - | 287.02 | 45.19 | 8.5e-3 | 20 |
| Col-0 | *35S:PLT1-GR* | DEX | 332.95 | 63.97 |  | 33 |
| *cle40-2* | *35S:PLT1-GR* | - | 173.11 | 33.90 | 8.3e-12 | 23 |
| *cle40-2* | *35S:PLT1-GR* | DEX | 338.68 | 69.00 |  | 20 |
| Col-0 | *35S:PLT2-GR* | - | 270.54 | 48.85 | 9.3e-7 | 17 |
| Col-0 | *35S:PLT2-GR* | DEX | 567.40 | 216.04 |  | 26 |
| *cle40-2* | *35S:PLT2-GR* | - | 200.08 | 38.73 | 1.4e-10 | 36 |
| *cle40-2* | *35S:PLT2-GR* | DEX | 442.43 | 161.21 |  | 25 |

| genotype | T-DNA | treatment | PRM cell number | standard deviation | *p*-value | n |
| --- | --- | --- | --- | --- | --- | --- |
| Col-0 |  | - | 44.33 | 5.10 | 0.58 | 12 |
| Col-0 |  | DEX | 44.06 | 4.47 |  | 18 |
| *cle40-2* |  | - | 27.22 | 3.42 | 0.14 | 27 |
| *cle40-2* |  | DEX | 29.00 | 3.91 |  | 19 |
| Col-0 | *35S:PLT1-GR* | - | 40.30 | 3.10 | 3.6e-13 | 20 |
| Col-0 | *35S:PLT1-GR* | DEX | 54.85 | 6.52 |  | 33 |
| *cle40-2* | *35S:PLT1-GR* | - | 24.78 | 5.18 | 1.1e-11 | 23 |
| *cle40-2* | *35S:PLT1-GR* | DEX | 50.70 | 10.36 |  | 20 |
| Col-0 | *35S:PLT2-GR* | - | 39.53 | 4.88 | 5.4e-4 | 17 |
| Col-0 | *35S:PLT2-GR* | DEX | 72.27 | 27.63 |  | 26 |
| *cle40-2* | *35S:PLT2-GR* | - | 26.26 | 5.33 | 2.9e-11 | 34 |
| *cle40-2* | *35S:PLT2-GR* | DEX | 62.64 | 22.78 |  | 25 |

**Table S4:**
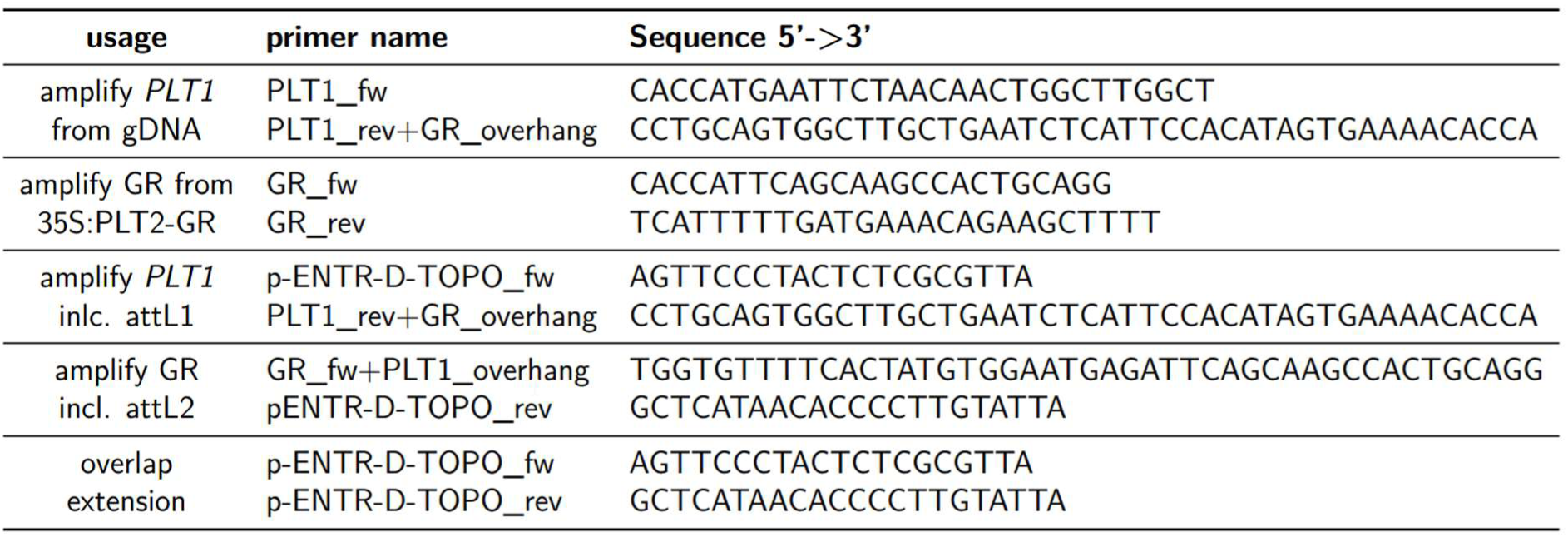
Primers used to generate *35S:PLT1-GR*.

